# Optimising biodiversity protection through artificial intelligence

**DOI:** 10.1101/2021.04.13.439752

**Authors:** Daniele Silvestro, Stefano Goria, Thomas Sterner, Alexandre Antonelli

**Affiliations:** Gothenburg Global Biodiversity Centre, Department of Biological and Environmental Sciences, University of Gothenburg, 405 30 Gothenburg, Sweden; Department of Biology, University of Fribourg, Switzerland and Swiss Institute of Bioinformatics; Thymia Limited, London, United Kingdom; Department of Economics, University of Gothenburg, Gothenburg, Sweden; Department of Plant Sciences, University of Oxford, South Parks Road, Oxford, OX1 3RB, United Kingdom; Royal Botanic Gardens, Kew, TW9 3AE, United Kingdom

**Keywords:** Artificial intelligence, Citizen science, Climate change, Human impact, Marxan, Remote sensing, Sustainable development, Systematic conservation planning

## Abstract

Over a million species face extinction, carrying with them untold options for food, medicine, fibre, shelter, ecological resilience, aesthetic and cultural values. There is therefore an urgent need to design conservation policies that maximise the protection of biodiversity and its contributions to people, within the constraints of limited budgets. Here we present a novel framework for spatial conservation prioritisation that combines simulation models, reinforcement learning and ground validation to identify optimal policies. Our methodology, CAPTAIN (Conservation Area Prioritisation Through Artificial Intelligence Networks), quantifies the trade-off between the costs and benefits of area and biodiversity protection, allowing the exploration of multiple biodiversity metrics. Under a fixed budget, our model protects substantially more species from extinction than the random or naively targeted protection of areas. CAPTAIN also outperforms the most widely used software for spatial conservation prioritisation (Marxan) in 97% of cases and reduces species loss by an average of 40% under simulations, besides yielding prioritisation maps at substantially higher spatial resolution using empirical data. We find that regular biodiversity monitoring, even if simple and with a degree of inaccuracy – characteristic of citizen science surveys – substantially improves biodiversity outcomes. Given the complexity of people–nature interactions and wealth of associated data, artificial intelligence holds great promise for improving the conservation of biological and ecosystem values in a rapidly changing and resource-limited world.

---

Biodiversity is the variety of all life on Earth, from genes through to populations, species, functions and ecosystems. Alongside its own intrinsic value and ecological roles, biodiversity provides us with clean water, pollination, building materials, clothing, food and medicine, among many other physical and cultural contributions that species make to ecosystem services and people’s lives^1,2^. The contradiction is that our endeavours to maximise short-term benefits are depleting biodiversity and threatening the life-sustaining foundations of modern societies in the long-run^3^ (Box 1). This can help explain why, despite the risks, we are living in an age of mass extinction^4,5^ and massive population declines of wild species^6^. One million species are currently at risk of disappearing^7^, including nearly 40% (or c. 140,000) of all plant species^8,9^, which could provide solutions to current and future challenges such as emerging diseases and climate-resilient crops^10-12^. The imperative to feed and house massively growing human populations – with an estimated 2.4 billion more people by 2050 – together with increasing disruptions from climate change, will put tremendous pressure on the world’s last remaining native ecosystems and the species they contain. Since not a single of the 20 Aichi biodiversity targets agreed by 196 nations for the period 2011–2020 has been fully met^13^, there is now an urgent need to design more realistic and effective policies for a sustainable future^14^ that help deliver the conservation targets under the post-2020 Global Biodiversity Framework (https://www.cbd.int/).

## Box 1

**Socio-economic valuation of biodiversity**

The field of economics sees nature as an asset, analogous to diversity in portfolio management^3,43^. It provides valuable insurance that may prove vital^44^. Risks are reduced if two assets are negatively correlated. Weitzman^45^ incorporates the idea of redundancy when species share genes, and shows that optimal policy under a budget constraint will involve strict prioritisation of some species.

The value of ecosystem services hinges on ease of substitution by man-made capital. In some respects, man can “replace” a coral reef by artificial structures. Fish still find protection and food, but the artificial reef does not fully “replace” nature. When nature is irreplaceable, we speak of strong (rather than weak) sustainability; meaning more care must be taken to protect natural capital assets^46^.

We see massive degradation of natural assets that could have yielded a high return if properly managed by effective owners or stewards. Institutional mechanisms (including secure property rights) are vital to protect nature^47^. Still, private incentives to preserve biodiversity (e.g. research and development in biodiversity-based solutions) remain weak^48^, and strong policies are needed^14^.

Both benefits and costs of conservation vary considerably across space. Protection is typically expensive for farmland or real estate close to cities. Some land is cheap but protection enforcement expensive precisely because of remoteness. Some areas may be valuable because they have high biodiversity or endemism. What the framework we developed here strives for is high total biodiversity benefits with low land and opportunity costs, at the same time as it reduces risks associated with climate change uncertainties^49^. Spatial targeting of sites that increase biodiversity protected per monetary unit is cost effective, and may also include recreational and other values^22,50^ that we hope to develop more in future model versions.

There have been several theoretical and practical frameworks underlying biological conservation since the 1960s^15^. The field was initially focused on the conservation of nature for itself, without human interference, but gradually incorporated the bidirectional links to people – recognising our ubiquitous influence on nature, and the multi-faceted contributions we derive from it^1,6,15^. Throughout this progress, a critical step has been the identification of priority areas for targeted protection, restoration planning and impact avoidance – triggering the development of the field of spatial conservation prioritisation or systematic conservation planning^16-18^. While humans and wild species are increasingly sharing the same space^19^, the preservation of largely intact nature remains critical for safeguarding many species and ecosystems, such as tropical rainforests.

Several tools and algorithms have been designed to facilitate systematic conservation planning^20^. They often allow the exploration and optimisation of trade-offs between variables, something not readily available in Geographic Information Systems^21^, which can lead to substantial economic, social and environmental gains^22^. While the initial focus has been on maximising the protection of species while minimising costs, additional parameters can sometimes be modelled, such as species rarity and threat, total protected area and evolutionary diversity^20,23,24^. The most widely used method so far, Marxan^25^ seeks to identify a set of protected areas that collectively allow particular conservation targets to be met under minimal costs, using a simulated annealing optimization algorithm. Despite its usefulness and popularity, Marxan and similar methods^20^ are designed to optimize a one-time policy, do not directly incorporate changes through time, and assume a single initial gathering of biodiversity and cost data (although temporal aspects can be explored by manually updating and re-running the models, under various targets^26^). In addition, the optimized conservation planning does not explicitly incorporate climate change, variation in anthropogenic pressure (although varying threat probabilities are dealt with in newer software extensions of Marxan^27,28^), or species-specific sensitivities to such changes.

Here we tackle the challenge of optimising biodiversity protection in a complex and rapidly evolving world by harnessing the power of artificial intelligence (AI). We develop a novel tool for systematic conservation planning (Fig. 1) to explore – through simulations and empirical analyses – multiple previously identified trade-offs in real-world conservation, and to evaluate the impact of data gathering on specific outcomes^29^. We also explore the impact of species-specific sensitivity to geographically varying local disturbances (e.g., as a consequence of new roads, mining, trawling or other forms of economic activity with negative impacts on natural ecosystems) and climate change (overall temperature increases, as well as short-term variations to reflect extreme weather events). We name our framework CAPTAIN (Conservation Area Prioritisation Through Artificial Intelligence Networks). Our framework is particularly applicable to sessile organisms, such as forest trees or reef corals, but implements mechanisms of offspring dispersal (e.g., seeds or larvae) that could be easily adaptable to many other systems.

**Fig. 1.**
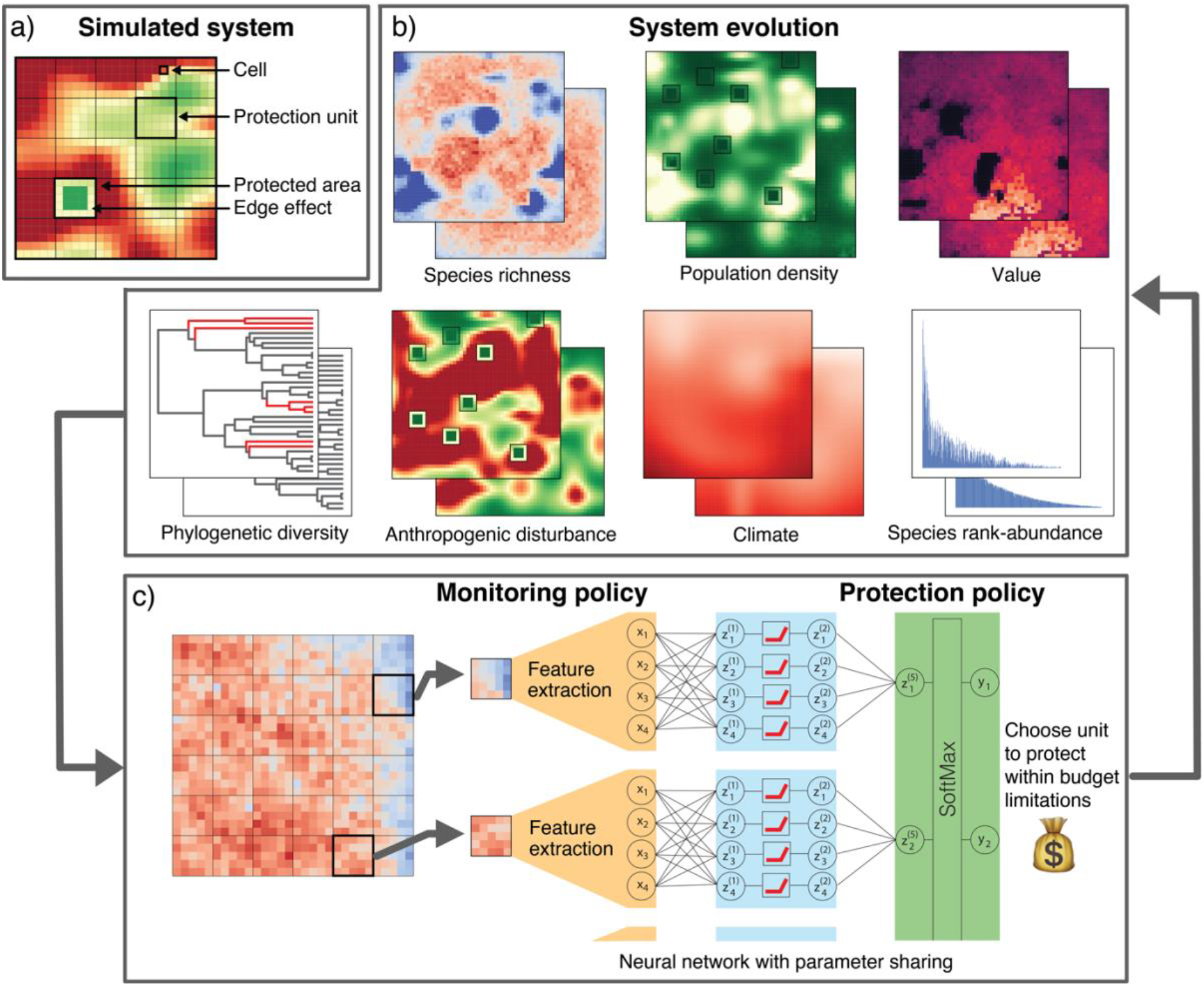
The CAPTAIN reinforcement learning framework. **a**, A simulated system – which could be the equivalent to a country state, an island or a large coral reef – consists of individual cells, each with a number of individuals of various species. Once a protection unit is identified and protected, its human-driven disturbance (e.g., forest loggings) will immediately reduce to an arbitrarily low level, except for the well-known edge effect^33^ characterised by intermediate levels of disturbance. All 44 model parameters are provided with initial default values but are fully customisable (see Tables S1–S2). Simulated systems evolve through time (b), which are used to optimize a conservation policy (c) and to evaluate its performance. In empirical applications of the CAPTAIN framework, the simulated system is replaced with available biodiversity and disturbance data. **b-c**, Analysis flowchart integrating simulations and AI modules to maximise selected outcomes (e.g., species richness). **b**, System evolution between two points in time, in relation to six variables: species richness, population density, economic value, phylogenetic diversity, anthropogenic disturbance, climate and species-rank abundance (see Animation S1 for a time-lapse video depicting these and additional attributes). Note that site protection does not protect species from climate change. **c**, Biodiversity features (species presence per cell at a minimum, plus their abundance under full monitoring schemes; see Methods and Box 2 for advances in data-gathering approaches) are extracted from the system at regular steps, which are then fed into a neural network that learns from the system’s evolution to identify conservation policies that maximise a reward, such as protection of the maximum species diversity within a fixed budget.

Within AI, we implement a reinforcement learning (RL) framework based on our spatially explicit simulation of biodiversity and its evolution through time in response to anthropogenic pressure and climate change. The RL algorithm is designed to find an optimal balance between data generation (learning from the current state of a system, also termed ‘exploration’) and action (called ‘exploitation’ – the effect of which is quantified by the outcome, also termed reward). It optimises a conservation policy that develops over time, thus being particularly suitable for designing policies and testing their short- and long-term effects. Actions are decided based on the state of the system through a neural network, whose parameters are estimated within the RL framework to maximise the reward. Although AI solutions have been previously proposed and to some extent are already used in conservation science^30,31^, to our knowledge RL has only been advocated^32^ but not yet implemented in conservation tools.

We combine these AI components with a module that allows us to maximise various biodiversity metrics (including species richness and phylogenetic diversity) within a given budget constraint (Fig. 1). We investigate through simulations the importance of data gathering, focusing on frequency (a single ‘Initial’, or multiple ‘Recurrent’ inputs of biodiversity data), level of detail (‘Simple’ species reports of species presence, or ‘Full’ monitoring of all species in targeted groups, and their population sizes) and accuracy (no errors in the recording, or allowing for a fraction of mistakes in species identifications characteristic of citizen science efforts). Our platform enables us to assess the influence of model assumptions on the reward, mimicking the use of counterfactual analyses^24^.

We use CAPTAIN to assess the following questions: i) What role does data gathering strategy play for effective conservation? ii) What trade-offs exist depending on the optimised variable, such as species richness, economic value or total area protected? iii) What can the simulation framework teach us in terms of winners and losers – i.e., which traits characterise the species and areas protected over time? and iv) How does our framework perform compared with the state-of-the-art model for conservation planning Marxan^25^? Finally, we demonstrate the usefulness of our framework to an empirical dataset of endemic trees of Madagascar.

## RESULTS AND DISCUSSION

### Impact of data gathering strategy

We find that Full Recurrent Monitoring (where the system is monitored at each time step, including species presence and abundance) results in the smallest species loss: it succeeds in protecting on average 35% more species than a random protection policy (Fig. 2a; Table S3). A very similar outcome (32% improvement) is generated by the Citizen science Recurrent Monitoring strategy (where only presence/absence of species are recorded in each cell) with a degree of error (Fig. 2b; see Methods). This is noteworthy because the information it requires, although not currently available at high spatial resolution for most regions and taxonomic groups, could to some extent be obtained through modern technologies, such as remote sensing and environmental DNA, and for accessible locations also citizen science^34^ in cost-efficient ways (Box 2).

**Fig. 2.**
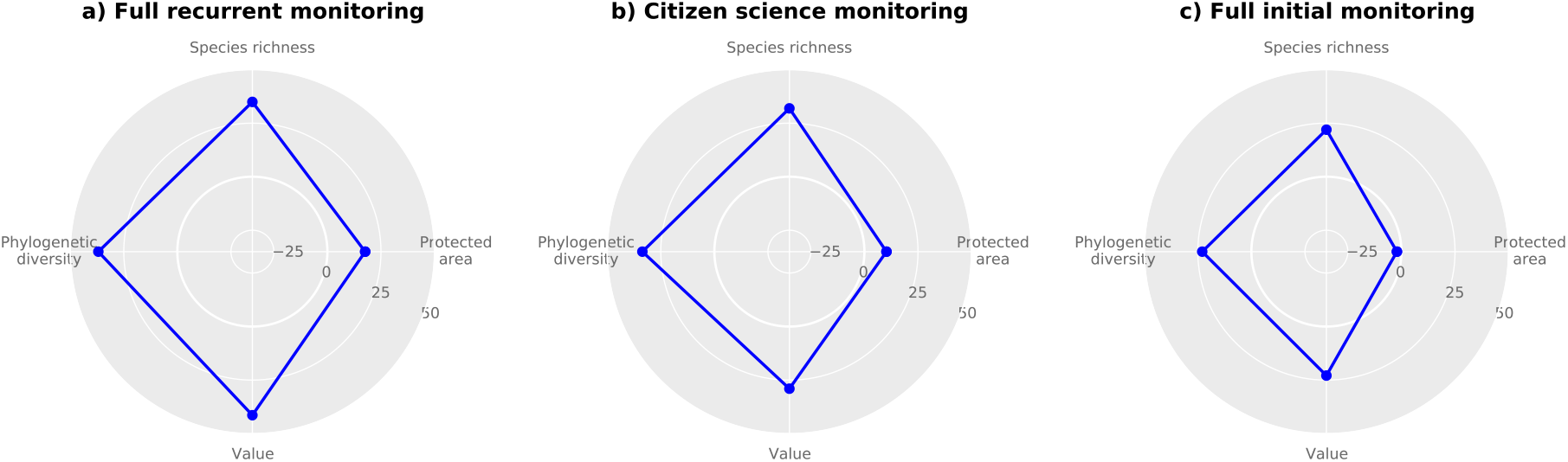
Impact of monitoring strategies on biodiversity protection. **a-c**, Outcome of policies designed to minimise species loss based on different monitoring strategies: **a**, Full Recurrent Monitoring (of species presence and abundance at each time step); **b**, Citizen science Recurrent Monitoring (limited to species presence/absence with some error at each time step); **c**, Full Initial Monitoring (species presence and abundance only at initial time). The blue polygons show the reduced loss of species, total protected area, accumulated species values and phylogenetic diversity between a random protection policy (grey shaded area) and AI-optimised simulations. All results are averaged across 250 simulations. Each simulation was based on the same budget and resolution of the protection units (5×5) but differed in their initial natural system (species distributions, abundances, tolerances, phylogenetic relationships) and in the dynamics of climate change and disturbance patterns.

#### Box 2

**New techniques for biodiversity monitoring**

Our simulations show that the regular monitoring of species is critical for prioritising areas for conservation under changing anthropogenic and climate pressure (Fig. 2). But given scarce resources and vast areas to monitor, how can this be most efficiently accomplished?

For centuries, biodiversity inventories have been carried out by biodiversity experts. This work has comprised the collection, recording and eventual identification of biological samples, such as fertile tree branches or whole animals – a time- and resource-consuming process, resulting in taxonomically and spatially biased and incomplete biodiversity data^51^. New approaches are now speeding up and popularising biodiversity monitoring, including **environmental DNA, remote sensing** and **citizen science**.

Rather than locating and recording each individual species directly, soil samples can be taken by non-specialists and their environmental DNA (eDNA) contents subsequently analysed. This approach reveals the biotic composition of whole communities, including plants, animals, fungi and bacteria. Standardised sampling also allows direct comparison of taxonomic and phylogenetic diversity across multiple sites^52^, helping to inform on presence and absence of species. Challenges remain in completing reference databases for matching sequences to names, and in separating local versus transported DNA from nearby areas.

Citizen science initiatives such as ‘bio-blitzes’ (an intense period of biological surveying, often by amateurs) can help fill up critical data gaps, reduce monitoring costs and increase local awareness and engagement for biodiversity. An impressive 3.5 billion smartphones are now in use around the world^53^, creating unseen possibilities of linking millions of people to nature through species logging, using platforms such as iNaturalist^54^ which increasingly use image-recognition software and geolocations for automated species identification. In our simulations, even identification errors of up to 20% during monitoring steps (a conservative rate for many organism groups, such as birds, mammals or trees) have only a marginal impact on the biodiversity outcomes (Fig. 3c). [Photo credits: a-b: A.A.; c: Justin Moat].

Finally, remote sensing technologies can rapidly scan and characterise relatively large areas. Using 3-D sensing technology, they now hold the promise of automated identification of species using multi-spectrum images^55^. The spectral signatures of species can be produced by combining on-the-ground (visible to NIR spectrophotometry of plant material and terrestrial LiDAR) with from-the-air (hyperspectral and LiDAR UAV survey) data. This allows the isolation of individual objects to record the presence and abundance of species. The technology can also provide measures of vegetation structure (informing on an areas’ functional diversity) and ecosystem services, such as carbon storage, soil water content, and other parameters that could be easily implemented in our model to help the valuation and ranking of areas for protection, besides greatly improving the quality of habitat data publicly available.

The two monitoring strategies above outperform a Simple Initial Monitoring with no error, which only saves from extinction an average of 22% more species than a random policy (Fig. 2c; Table S3). This is because the policy ignores the temporal dynamics caused by disturbances, population and range changes, and the costs of area protection – all of which are likely to change through time in real-world situations. Since current methodologies for systematic conservation planning are static – relying on a similar initial data gathering as modelled here – this means their recommendations for area protection may be less reliable.

Our results differ from previous suggestions that area protection should not await prolonged data gathering^29^. This discrepancy can be explained by the nature of changes over time: while some systems may remain largely static over decades (e.g., tree species in old-growth forests), where a Simple Initial Monitoring could suffice, others may change drastically (e.g., alpine meadows or shallow-sea communities, where species shift their ranges rapidly in response to climatic and anthropogenic pressures); all such parameters can be tuned in our model.

To thoroughly explore the parameter space, each simulated system was initialised with different species composition and distributions and different anthropogenic pressure and climate change patterns (Figs. S1–S4). Because of this stochasticity, the reliability of the protection policies in relation to species loss varied across simulations. The policy based on Full Recurrent Monitoring was the most reliable, outperforming the baseline random policy in 99% of the simulations, while the Citizen science Recurrent Monitoring outperformed the random baseline in 92% of cases (Table S3). Both those policies are more reliable than the Full Initial Monitoring, which in addition to protecting fewer species on average (Fig. 2) also results in a slightly lower reliability of the outcome, outperforming the random policy in 89% of the simulations.

### Optimisation trade-offs

The policy objective, which determines the optimality criterion in our RL framework, significantly influences the outcome of the simulations. A policy maximizing species commercial values (such as timber price) tends to sacrifice more species in order to prioritise the protection of fewer, highly valuable ones. Under this scenario, species losses decrease by only 2.5% compared with the random baseline, while cumulative value losses decrease by about 70% (Table S3). Thus, a policy targeting exclusively the preservation of species with high economic value may have a strongly negative impact on the total protected species richness, phylogenetic diversity and even amount of protected area, compared with a policy minimizing species loss (Fig. 3a).

**Fig. 3.**
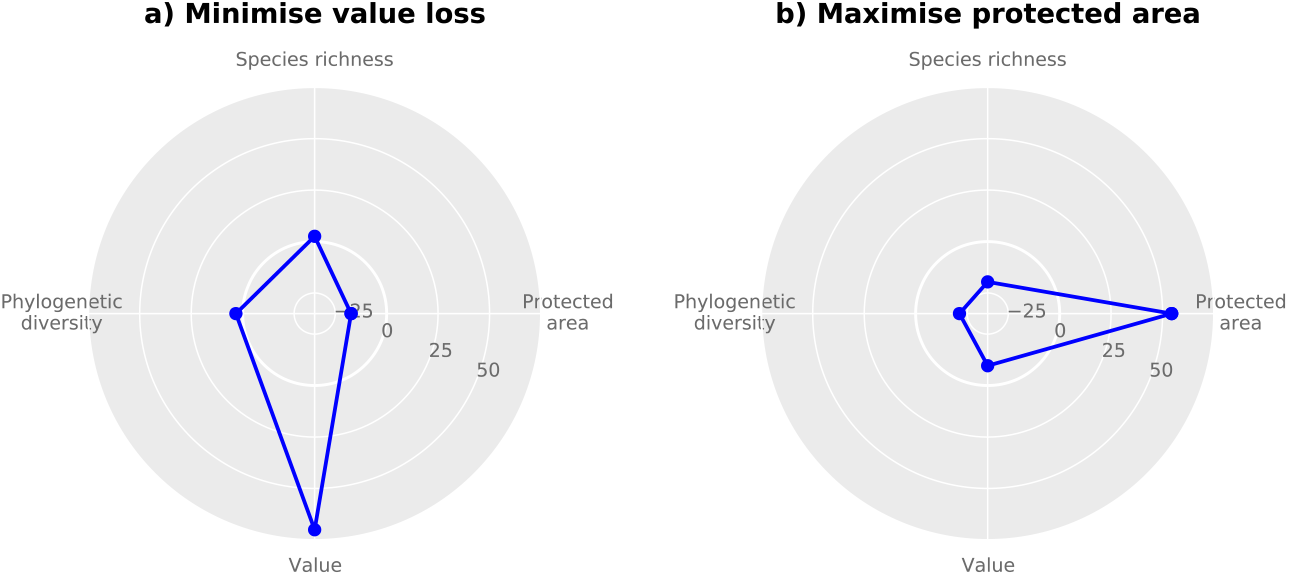
Trade-offs in conservation outcomes in relation to policy objectives. The plots show the outcome (averaged across 250 simulations) of different policy objectives based on Full Recurrent Monitoring. The policies were designed to (**a)** minimize value loss and (**b)** maximize the amount of protected area. The radial axis shows the percentage change compared with the baseline random policy, while the dashed grey polygons show the outcome of a policy with full recurrent monitoring optimized to minimize biodiversity loss (Fig. 2a).

A policy that maximises protected area results in a 54% increase in number of protected cells, by selecting those cheapest to buy; but it leads to substantial losses in species numbers, value and phylogenetic diversity, which are considerably worse than the random baseline, by resulting in 20% more species losses on average (Table S3). The decreased performance in terms of preventing extinctions is even more pronounced when compared with a policy minimizing species loss (Fig. 3b). This is an important finding, given that total protected area has been at the core of previous international targets for biodiversity (such as Aichi; https://www.cbd.int/sp/targets), and remains a key focus under the new post-2020 Global Biodiversity Framework under the Convention on Biological Diversity. Focusing on quantity (area protected) rather than quality (actual biodiversity protected) could inadvertently support political pressure for ‘residual’ reservation^35,36^ – the selection of new protected areas on land and at sea that are unsuitable for extractive activities, which may reduce costs and risks for conflicts but are likely suboptimal for biodiversity conservation.

As expected, the reliability for optimisations on economic value and total protected area was high for the respective policy objectives, decreasing the value loss in 96% of the simulations compared to the random baseline and increasing the amount of protected area in 100% of the simulations (Table S3). However, these policies resulted in highly inconsistent outcomes in terms of preventing species extinctions, with biodiversity losses not significantly different from those of the random baseline policy (Table S3). This indicates that economic value and total protected area should not be used as surrogates for biodiversity protection.

### Winners and losers

Focusing on the policy developed under Full Recurrent Monitoring and optimised on reducing species loss, we explored the properties of species that survived in comparison with those that went extinct, despite optimal area protection. Species that went extinct were characterised by relatively small initial range, small populations and intermediate or low resilience to disturbance (Fig. 4a). In contrast, species that survived had either low resilience but widespread ranges and high population sizes, or high resilience with small ranges and population sizes.

**Fig. 4.**
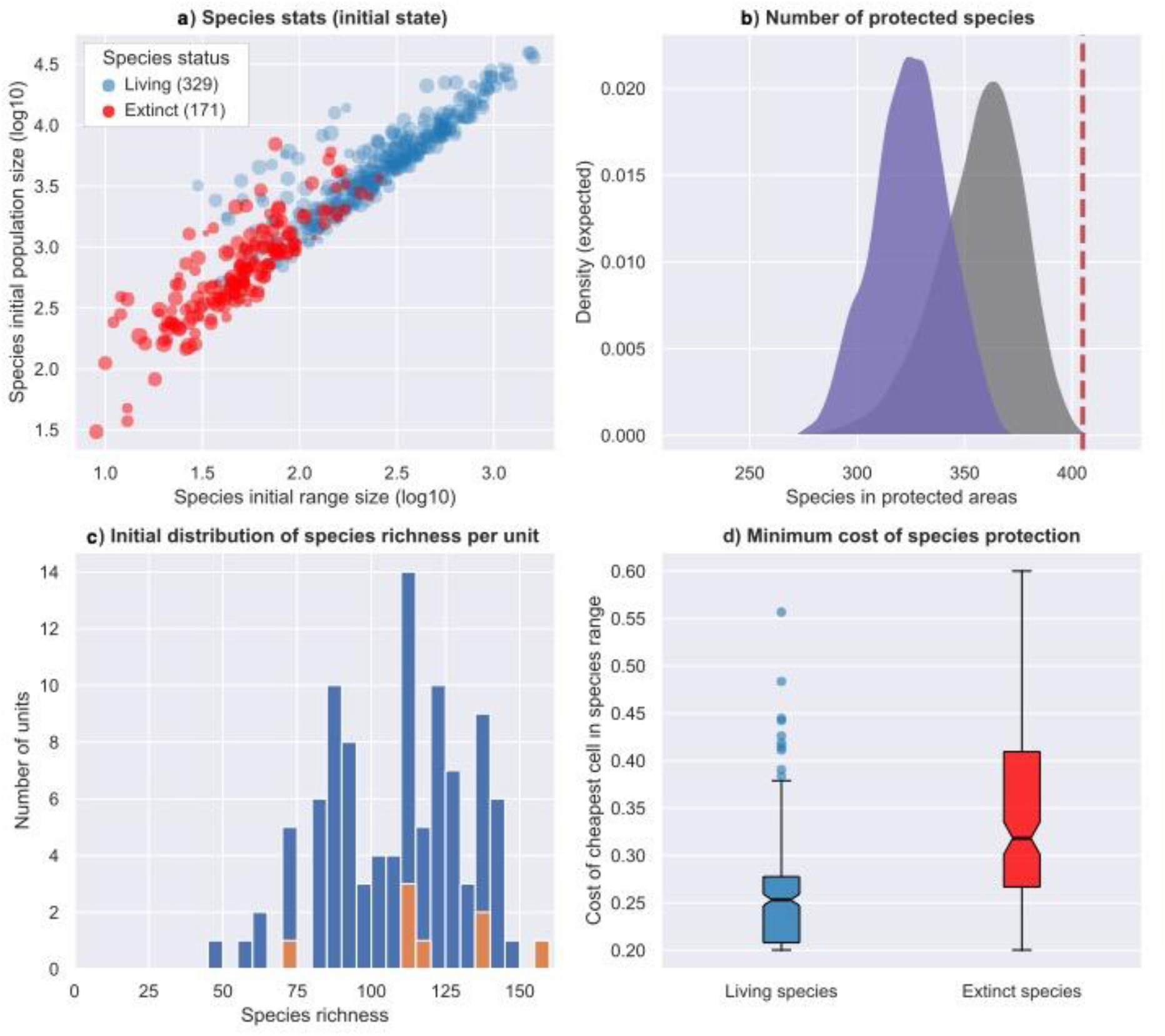
Summary statistics for one simulation optimised to reduce species loss and informed through Full Recurrent Monitoring. **a**, Living or surviving (blue circles) and locally extinct (red circles) species after a simulation of 30 time-steps with increasing disturbance and climate change. The X and Y axes show the initial range size and population size of the species (log10 transformed) and the size of the circles is proportional to the resilience of each species to anthropogenic disturbance and climate change, with smaller circles representing more sensitive species. **b**, Cumulative number of species encompassed in the eight protected units (5×5 cells) selected based on a policy optimised to minimise species loss. The grey density plot shows the expected distribution from 10,000 random draws, the purple shaded area shows the expected distribution when protected units are selected ‘naively’ (here, chosen among the top 20 most diverse ones), while the dashed red line indicates the number of species included in the units selected by the optimised CAPTAIN policy, which is higher than in 99.9% of the random draws. The optimised policy learned to maximise the total number of species included in protected units, thus accounting for their complementarity. Note that fewer species survived (329) in this simulation compared to how many were included in protected areas (405). This discrepancy is due to the effect of climate change, in which area protection does not play a role (Animation S1). Connectivity of protected areas could facilitate climate-induced range shifts. **c**, Species richness across the 100 protection units included in the area (blue), eight of which were selected to be protected (orange). The plot shows that the protection policy does not exclusively target units with the highest diversity. **d**, Distribution of the minimum cost of protection for living and extinct species. The plot shows the cost of the cell with the lowest cost of protection (as quantified at the last time step) within the initial range of each species.

We further assessed what characterised the grid cells that were selected for protection by the optimised policy. The cumulative number of species included in these cells was significantly higher than the cumulative species richness across a random set of cells of equal area (Fig. 4b). Thus, the model learns to protect a diversity of species assemblages to minimise species loss. Interestingly, the cells selected for protection did not include only areas with the highest species richness (Fig. 4c). This is important because areas with high richness constitute a ‘naive’ conservation target. Instead, protected cells spanned a range of areas with intermediate to high species richness, which reflects known differences among ecosystems or across environmental gradients and is more likely to increase protection complementarity for multiple species, a key factor incorporated by our software and some others^16,25,37^. Finally, extinct species tend to occur in more expensive regions within the system (Fig. 4d), corresponding to areas with high disturbance, which in turn might explain why they could not be effectively protected within the limited budget of the simulation.

If we instead examine which areas were protected under a policy maximizing total protected area, we find that they often include adjacent cells, for instance areas in a remote region, far from disturbance and therefore cheaper to buy and protect – typical of many current residual reserves, although less so for some newer conservation schemes^35,36^.

### Empirical applications and benchmarking

We evaluated our simulation framework by comparing its performance in optimizing policies against the current state-of-the-art tool for conservation prioritisation, Marxan^25^. We tested two monitoring policies: the Citizen science Recurrent and the Full Initial Monitoring, for which we could run analogous models within Marxan (see Methods). The analysis of 250 simulations showed that biodiversity loss was smallest in simulations using CAPTAIN with recurrent monitoring. Specifically, species loss using Marxan optimization with recurrent monitoring was on average 24% higher than using the CAPTAIN model (standard deviation, SD =18.9%) and resulted in a higher number of extinctions in 92% of the simulations (Fig. 5). Our methodology was even more successful when applying a one-time monitoring policy, which is the most common approach for identifying and delimiting protected areas worldwide; the CAPTAIN model outperformed Marxan in 96.8% of the simulations, with the latter resulting in a 40% average increase of species loss (SD = 26.7%). We interpret part of this difference in performance as the result of a different optimization objective, whereby Marxan seeks to find the cheapest distribution of protected units that meet a conservation target (e.g., protecting a minimum fraction of all species’ initial range), without explicitly incorporating a finite budget as a limiting factor.

**Fig. 5.**
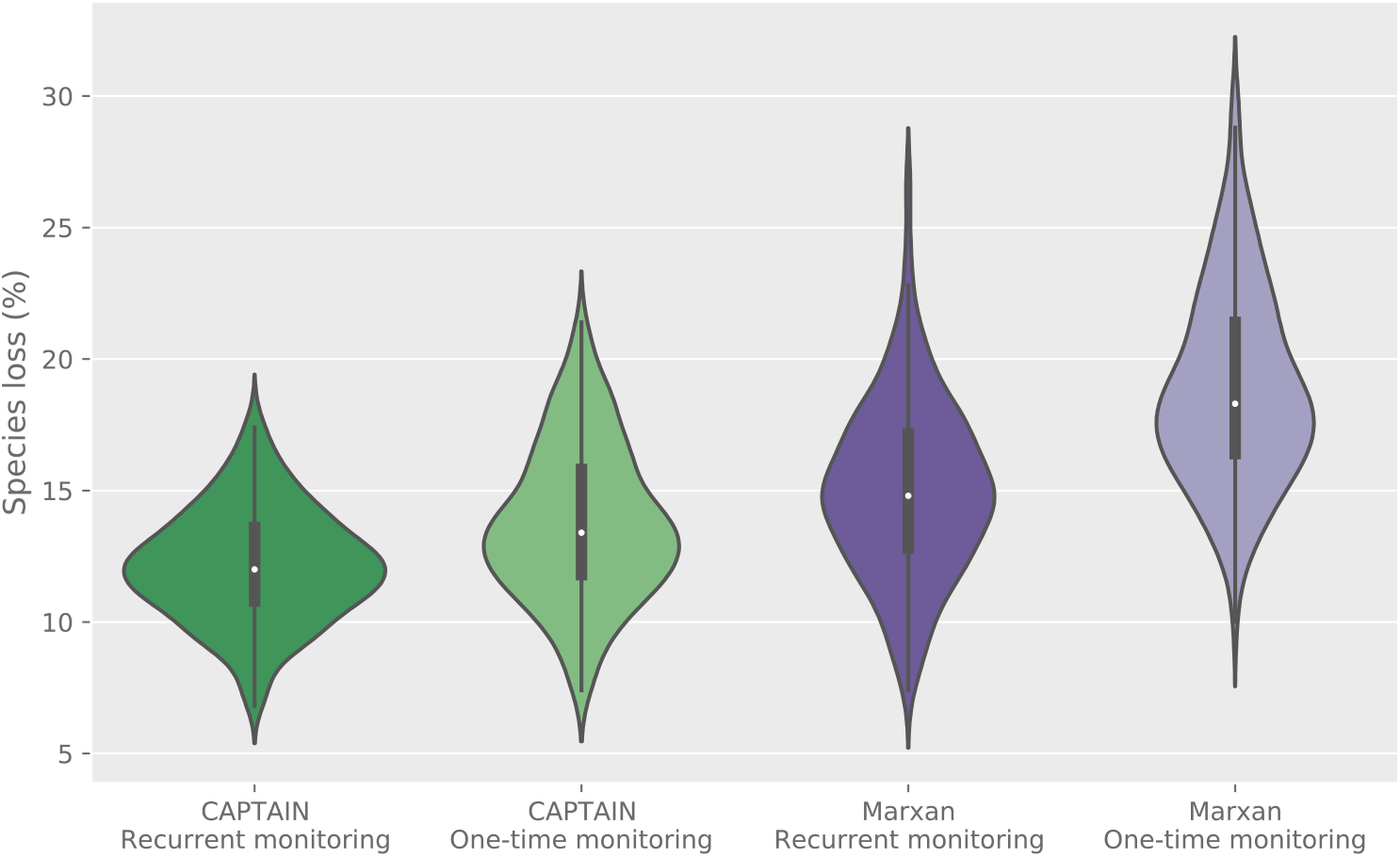
Benchmarking of CAPTAIN through simulations. The violin plots show the distribution of species loss outcomes across 250 simulations run under different monitoring policies based on the CAPTAIN framework developed here and on Marxan^25^. Species loss is expressed as a percentage of the total initial number of species in the simulated systems. See text for details of the analyses.

To demonstrate the potential applicability of our framework and its scalability to large datasets, we analysed a Madagascar biodiversity dataset recently used in a systematic conservation planning experiment^38^ using Marxan^25^. The dataset included 22,394 protection units (5 x 5 km) and presence-absence data for 1,517 endemic tree species (see Methods for more details). Our results, here constrained to protect the same number of units as in the Marxan analysis, showed some overlap with the Marxan output for the highest priority areas, such as in the north (∼ 13º S, 49º E), northeast (∼ 15º S, 50º E) and centre (∼ 20º S, 47º E) of the country (Figs. 6, S5–S6). Protection units with a Marxan priority greater than 75% were also assigned relatively high priority in CAPTAIN (on average 68%), while units with Marxan rank below 25% had low CAPTAIN rank (< 17%). This indicates a degree of consistency in the ranking of protection units across different methods. However, CAPTAIN additionally identified several other high priority areas across the country, within a wide region ranked by Marxan with only uniform average priority (ranked 25-75%). Importantly, CAPTAIN was able to identify priority areas for conservation at very high spatial resolution (Fig. 6b), making it a useful tool for informing on-the-ground decisions by landowners and policymakers.

**Fig. 6.**
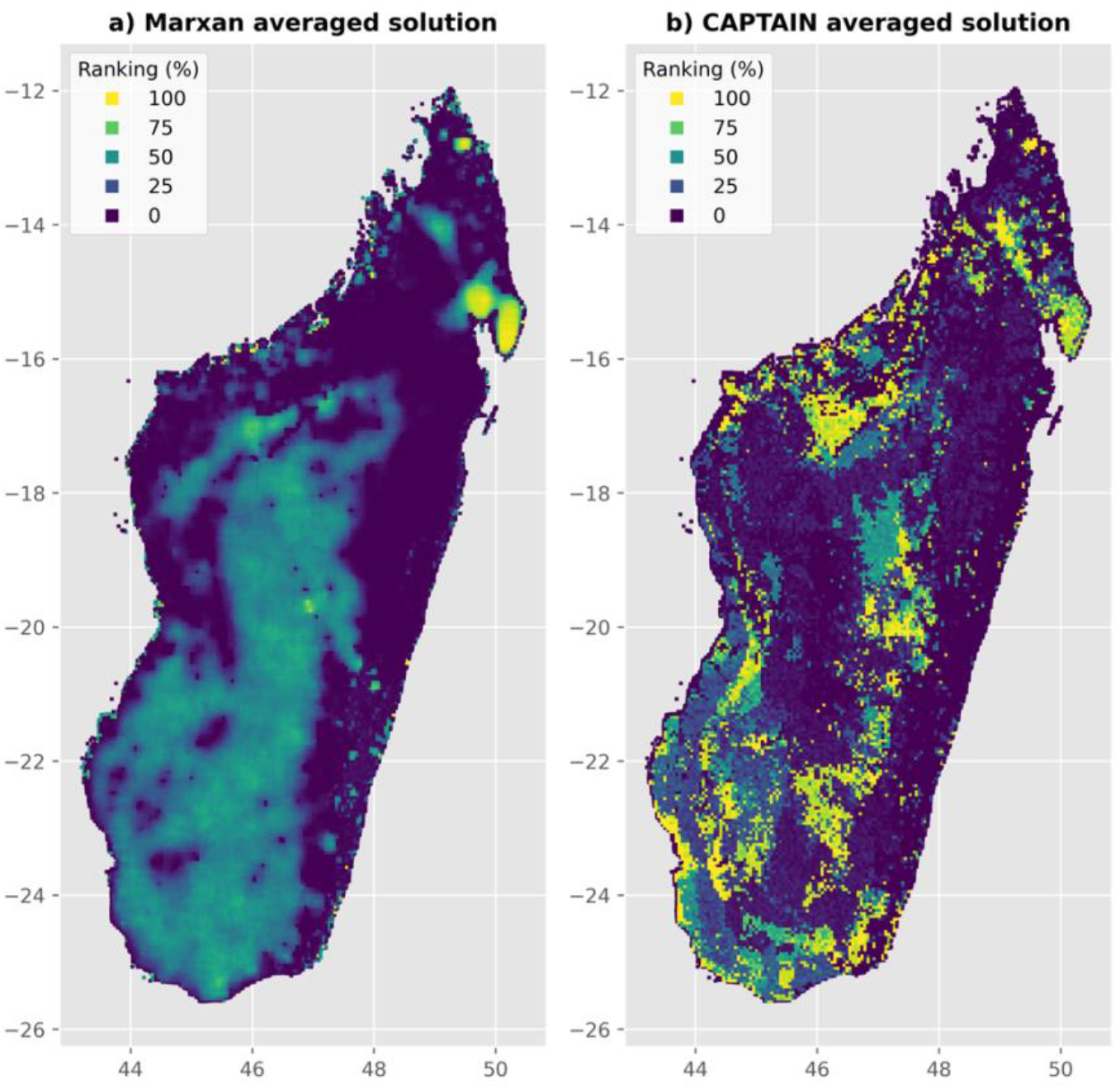
Empirical validation of CAPTAIN. The maps show a ranking of priority areas for protection across Madagascar based on the distribution of endemic trees. **a)**, protection unit ranks resulting from an analysis using Marxan^25^ (as in Fig. S7d of Carrasco et al.^38^); **b)**, equivalent CAPTAIN results based on a Citizen science Recurrent Monitoring policy.

## Conclusions

In contrast to many short-lived decisions by governments, the selection of which areas in a country’s territory should be protected will have long-term repercussions. Protecting the right areas will help safeguard natural assets and their contributions for the future. Choosing suboptimal areas, by contrast, will lead to the loss of species, phylogenetic diversity, socio-economic value and ecological functions. As public funding for nature conservation is scarce and needs to compete with many other important environmental issues such as climate change or the spread of toxins, there is little room for mistakes.

The framework presented here provides a spatially explicit way to explore the complex decisions facing policymakers and conservationists today, focusing on data gathering, placement of protected areas and policy objective – all within the context of a rapidly changing world affected by local anthropogenic disturbances and large-scale climate change. Our results highlight the importance of regularly gathering basic biodiversity information and demonstrate substantial trade-offs in biodiversity outcome depending on the policy being optimised – such as maximizing the protection of species or areas.

As the number of standardised high-resolution biological datasets is increasing (e.g.,^39^), supported by the use of new and cost-effective monitoring technologies (Box 2), our approach offers new opportunities for research and conservation. The flexibility of our model, offered by its nearly 40 variables (see Methods), could be easily expanded and adapted to almost any empirical dataset. Future implementations could model additional variables, such as functional diversity and other biodiversity and socio-economic metrics.

Artificial intelligence techniques should not replace human judgment, and ultimately investment decisions will be based on more than just the simple parameters implemented in our models, including careful consideration of people’s manifold interactions with nature^1,15^. More sophisticated measures of value could be incorporated in future work. It is also crucial to recognise the importance of ensuring the right conditions required for effective conservation of protected areas in the long term^40,41^. However, it is now time to acknowledge that the sheer complexity of biological systems, multiplied by the increasing disturbances in a changing world, cannot be fully grasped by the human mind. As we progress in what many are calling the most decisive decade for biodiversity^6,42^, we must take advantage of powerful tools that help us steward the planet’s remaining ecosystems in sustainable ways – for the benefit of people and all life on Earth.

## REFERENCES

1 Díaz, S. et al. Assessing nature’s contributions to people. Science 359, 270–272, doi:10.1126/science.aap8826 (2018).

2 Chaplin-Kramer, R. et al. Global modeling of nature’s contributions to people. Science 366, 255–258, doi:10.1126/science.aaw3372 (2019).

3 Dasgupta, P. The Economics of Biodiversity: The Dasgupta Review. London: HM Treasury (2021).

4 Barnosky, A. D. et al. Has the Earth’s sixth mass extinction already arrived? Nature 471, 51–57, doi:10.1038/nature09678 (2011).

5 Andermann, T., Faurby, S., Turvey, S. T., Antonelli, A. & Silvestro, D. The past and future human impact on mammalian diversity. Science Advances 6, eabb2313, doi:10.1126/sciadv.abb2313 (2020).

6 Leclère, D. et al. Bending the curve of terrestrial biodiversity needs an integrated strategy. Nature 585, 551–556, doi:10.1038/s41586-020-2705-y (2020).

7 IPBES. Global Assessment on Biodiversity and Ecosystem Services. (2019).

8 Nic Lughadha, E. et al. Extinction risk and threats to plants and fungi. PLANTS, PEOPLE, PLANET 2, 389–408, doi:10.1002/ppp3.10146 (2020).

9 Antonelli, A. et al. State of the World’s Plants and Fungi 2020. (Royal Botanic Gardens, Kew, 2020).

10 Howes, M.-J. R. et al. Molecules from nature: Reconciling biodiversity conservation and global healthcare imperatives for sustainable use of medicinal plants and fungi. PLANTS, PEOPLE, PLANET 2, 463–481, doi:10.1002/ppp3.10138 (2020).

11 Kersey, P. J. et al. Selecting for useful properties of plants and fungi – Novel approaches, opportunities, and challenges. PLANTS, PEOPLE, PLANET 2, 409–420, doi:10.1002/ppp3.10136 (2020).

12 Seddon, N. et al. Understanding the value and limits of nature-based solutions to climate change and other global challenges. Philos. Trans. R. Soc. Lond. B Biol. Sci. 375, 20190120, doi:doi:10.1098/rstb.2019.0120 (2020).

13 Diversity, S. o. t. C. o. B. Global Biodiversity Outlook 5 – Summary for Policy Makers. (2020).

14 Sterner, T. et al. Policy design for the Anthropocene. Nature Sustainability 2, 14–21, doi:10.1038/s41893-018-0194-x (2019).

15 Mace, G. M. Whose conservation? Science 345, 1558–1560, doi:10.1126/science.1254704 (2014).

16 Moilanen, A. Generalized Complementarity and Mapping of the Concepts of Systematic Conservation Planning. Conservation biology : the journal of the Society for Conservation Biology 22, 1655–1658, doi:10.1111/j.1523-1739.2008.01043.x (2008).

17 Margules, C. & Sarkar, S. Systematic conservation planning. (Cambridge University Press, 2007).

18 Margules, C. R. & Pressey, R. L. Systematic conservation planning. Nature 405, 243–253, doi:10.1038/35012251 (2000).

19 Obura, D. O. et al. Integrate biodiversity targets from local to global levels. Science 373, 746–748, doi:10.1126/science.abh2234 (2021).

20 Moilanen, A., Wilson, K. & Possingham, H. Spatial conservation prioritization: quantitative methods and computational tools. (Oxford University Press, 2009).

21 Honeck, E., Sanguet, A., Schlaepfer, M. A., Wyler, N. & Lehmann, A. Methods for identifying green infrastructure. SN Applied Sciences 2, 1916, doi:10.1007/s42452-020-03575-4 (2020).

22 Bateman, I. J. et al. Bringing Ecosystem Services into Economic Decision-Making: Land Use in the United Kingdom. Science 341, 45–50, doi:10.1126/science.1234379 (2013).

23 Carvalho, S. et al. Spatial conservation prioritization of biodiversity spanning the evolutionary continuum. Nature Ecology & Evolution 1, 0151, doi:10.1038/s41559-017-0151 (2017).

24 Sacre, E., Weeks, R., Bode, M. & Pressey, R. L. The relative conservation impact of strategies that prioritize biodiversity representation, threats, and protection costs. Conservation Science and Practice 2, e221, doi:https://doi.org/10.1111/csp2.221 (2020).

25 Ball, I. R., Possingham, H. P. & Watts, M. Marxan and relatives: software for spatial conservation prioritisation. Spatial conservation prioritisation: Quantitative methods and computational tools, 185–195 (2009).

26 Pressey, R. L., Mills, M., Weeks, R. & Day, J. C. The plan of the day: Managing the dynamic transition from regional conservation designs to local conservation actions. Biol. Conserv. 166, 155–169, doi:https://doi.org/10.1016/j.biocon.2013.06.025 (2013).

27 Watts, M., Klein, C. J., Tulloch, V. J. D., Carvalho, S. B. & Possingham, H. P. Software for prioritizing conservation actions based on probabilistic information. Conserv. Biol. 35, 1299–1308, doi:https://doi.org/10.1111/cobi.13681 (2021).

28 Tulloch, V. J. et al. Incorporating uncertainty associated with habitat data in marine reserve design. Biol. Conserv. 162, 41–51, doi:https://doi.org/10.1016/j.biocon.2013.03.003 (2013).

29 Grantham, H. S., Wilson, K. A., Moilanen, A., Rebelo, T. & Possingham, H. P. Delaying conservation actions for improved knowledge: how long should we wait? Ecol. Lett. 12, 293–301, doi:https://doi.org/10.1111/j.1461-0248.2009.01287.x (2009).

30 Zizka, A., Silvestro, D., Vitt, P. & Knight, T. M. Automated conservation assessment of the orchid family with deep learning. Conserv. Biol. n/a, doi:https://doi.org/10.1111/cobi.13616 (2020).

31 Gomes, C. et al. Computational sustainability: computing for a better world and a sustainable future. Commun. ACM 62, 56–65, doi:10.1145/3339399 (2019).

32 Spring, D. A., Cacho, O., Mac Nally, R. & Sabbadin, R. Pre-emptive conservation versus “fire-fighting”: A decision theoretic approach. Biol. Conserv. 136, 531–540, doi:https://doi.org/10.1016/j.biocon.2006.12.024 (2007).

33 Murcia, C. Edge effects in fragmented forests: implications for conservation. Trends Ecol. Evol. 10, 58–62 (1995).

34 Kosmala, M., Wiggins, A., Swanson, A. & Simmons, B. Assessing data quality in citizen science. Front. Ecol. Environ. 14, 551–560, doi:https://doi.org/10.1002/fee.1436 (2016).

35 Devillers, R. et al. Reinventing residual reserves in the sea: are we favouring ease of establishment over need for protection? Aquat. Conserv.: Mar. Freshwat. Ecosyst. 25, 480–504, doi:https://doi.org/10.1002/aqc.2445 (2015).

36 Vieira, R. R. S., Pressey, R. L. & Loyola, R. The residual nature of protected areas in Brazil. Biol. Conserv. 233, 152–161, doi:https://doi.org/10.1016/j.biocon.2019.02.010 (2019).

37 Dilkina, B. et al. Trade-offs and efficiencies in optimal budget-constrained multispecies corridor networks. Conserv. Biol. 31, 192–202 (2017).

38 Carrasco, J., Price, V., Tulloch, V. & Mills, M. Selecting priority areas for the conservation of endemic trees species and their ecosystems in Madagascar considering both conservation value and vulnerability to human pressure. Biodivers. Conserv. 29, 1841–1854 (2020).

39 Bruelheide, H. et al. sPlot – A new tool for global vegetation analyses. Journal of Vegetation Science 30, 161–186, doi:10.1111/jvs.12710 (2019).

40 Geldmann, J., Manica, A., Burgess, N. D., Coad, L. & Balmford, A. A global-level assessment of the effectiveness of protected areas at resisting anthropogenic pressures. Proc. Natl. Acad. Sci. USA 116, 23209–23215, doi:10.1073/pnas.1908221116 (2019).

41 Geldmann, J. et al. A global analysis of management capacity and ecological outcomes in terrestrial protected areas. Conservation Letters 11, e12434, doi:https://doi.org/10.1111/conl.12434 (2018).

42 Díaz, S. et al. Set ambitious goals for biodiversity and sustainability. Science 370, 411–413, doi:10.1126/science.abe1530 (2020).

43 Brock, W. A. & Xepapadeas, A. Valuing Biodiversity from an Economic Perspective: A Unified Economic, Ecological, and Genetic Approach. Am Econ Rev 93, 1597–1614, doi:10.1257/000282803322655464 (2003).

44 Goeschl, T. & Swanson, T. The social value of biodiversity for R&D. Environmental and Resource Economics 22, 477–504 (2002).

45 Weitzman, M. L. The Noah’s ark problem. Econometrica, 1279-1298 (1998).

46 Barbier, E. B. The concept of natural capital. Oxford Review of Economic Policy 35, 14–36, doi:10.1093/oxrep/gry028 (2019).

47 Dasgupta, P. The Dasgupta Review –Independent Review on the Economics of Biodiversity. Interim Report. (2020).

48 Simpson, R. D., Sedjo, R. A. & Reid, J. W. Valuing biodiversity for use in pharmaceutical research. Journal of Political Economy 104, 163–185 (1996).

49 Ando, A. W. & Mallory, M. L. Optimal portfolio design to reduce climate-related conservation uncertainty in the Prairie Pothole Region. Proc. Natl. Acad. Sci. USA 109, 6484–6489, doi:10.1073/pnas.1114653109 (2012).

50 Bateman, I. J. et al. Conserving tropical biodiversity via market forces and spatial targeting. Proceedings of the National Academy of Sciences 112, 7408–7413 (2015).

51 Meyer, C., Weigelt, P. & Kreft, H. Multidimensional biases, gaps and uncertainties in global plant occurrence information. Ecology Letters 19, 992–1006, doi:10.1111/ele.12624 (2016).

52 Ritter, C. D. et al. The pitfalls of biodiversity proxies: Differences in richness patterns of birds, trees and understudied diversity across Amazonia. Scientific reports 9, 19205, doi:10.1038/s41598-019-55490-3 (2019).

53 Deyan, G. 61+ Revealing Smartphone Statistics For 2020. (2019). <https://techjury.net/stats-about/smartphone-usage/#gref>.

54 Wäldchen, J., Rzanny, M., Seeland, M. & Mäder, P. Automated plant species identification—Trends and future directions. PLoS Comp. Biol. 14, e1005993, doi:10.1371/journal.pcbi.1005993 (2018).

55 Baena, S., Moat, J., Whaley, O. & Boyd, D. S. Identifying species from the air: UAVs and the very high resolution challenge for plant conservation. PLOS ONE 12, e0188714, doi:10.1371/journal.pone.0188714 (2017).

56 Barto, A. G. Reinforcement Learning and Dynamic Programming. IFAC Proceedings Volumes 28, 407–412, doi:https://doi.org/10.1016/S1474-6670(17)45266-9 (1995).

57 Weng, L. 2020).

58 Salimans, T., Ho, J., Chen, X., Sidor, S. & Sutskever, I. Evolution strategies as a scalable alternative to reinforcement learning. arXiv preprint 1703.03864 (2017).

59 Sutton, R. S., McAllester, D. A., Singh, S. P. & Mansour, Y. in NIPs. 1057-1063 (Citeseer).

60 ter Steege, H. et al. Hyperdominance in the Amazonian Tree Flora. Science 342, doi:10.1126/science.1243092 (2013).

61 Stadler, T. Simulating trees with a fixed number of extant species. Syst. Biol. 60, 676–684 (2011).

62 Game, E. T. & Grantham, H. S. Marxan user manual. For Marxan version 1 (2008).

63 Tulloch, V. J. et al. Incorporating uncertainty associated with habitat data in marine reserve design. Biol. Conserv. 162, 41–51 (2013).

